# Astrocytic ablation of truncated TrkB receptors reduces amyloid plaques and prevents synaptic loss in 5xFAD mice

**DOI:** 10.64898/2026.10.20.760443

**Authors:** Robert Bass, Hannah V. Masterson, Yuting Chen, Baoji Xu

## Abstract

Brain-derived neurotrophic factor (BDNF) is essential for nervous system development and function and exerts its effects primarily through the tropomyosin receptor kinase B (TrkB). The *Ntrk2* gene encodes a full-length receptor (TrkB.FL) and two truncated isoforms (TrkB.T1 and TrkB.T2, collectively TrkB.T) that lack the intracellular tyrosine kinase domain. TrkB.T is expressed predominantly in astrocytes and is thought to inhibit BDNF-TrkB.FL signaling by sequestering BDNF and forming signaling-deficient heterodimers with TrkB.FL. Here, we investigated whether selective ablation of astrocytic TrkB.T could ameliorate Alzheimer’s disease (AD) pathology by enhancing endogenous BDNF-TrkB.FL signaling. Astrocyte-specific deletion of the *Ntrk2* gene at 5 weeks of age reduced cortical TrkB.T expression by more than 80% without affecting TrkB.FL levels. TrkB.T ablation increased the size of Bassoon-positive presynaptic active zones, consistent with enhanced BDNF-TrkB.FL signaling. Remarkably, astrocytic TrkB.T ablation reduced β-amyloid plaque burden by more than 60% in the cortex, hippocampus, and subiculum of 6-month-old 5xFAD mice, despite having no detectable effect on microgliosis or astrogliosis. Furthermore, TrkB.T ablation prevented reductions in the density and size of PSD-95-positive postsynaptic sites in the cortex and hippocampus of 5xFAD mice at 1 year old. Together, these findings identify astrocytic TrkB.T as an important regulator of amyloid pathology and synaptic integrity in an AD mouse model. Our results suggest that selectively reducing TrkB.T expression represents a promising therapeutic strategy to enhance endogenous neuroprotective BDNF-TrkB.FL signaling, reduce amyloid pathology, and preserve synaptic integrity in AD.

**Significance Statement:** Alzheimer’s disease (AD) is characterized by progressive synaptic loss and neuronal degeneration, yet effective therapies that preserve synaptic integrity remain limited. Brain-derived neurotrophic factor (BDNF) signaling through its receptor TrkB promotes neuronal survival and synaptic plasticity, but this neuroprotective pathway is impaired in AD. Here, we demonstrate that selective ablation of truncated TrkB receptors in astrocytes markedly reduces β-amyloid plaque burden and prevents synaptic loss in 5XFAD mice without altering microgliosis or astrogliosis. These findings reveal an unexpected role for astrocytic truncated TrkB receptors in regulating amyloid pathology and synaptic integrity. Our study identifies astrocytic truncated TrkB receptors as promising therapeutic targets for AD and suggests that removing an endogenous inhibitor of BDNF signaling may provide a novel neuroprotective strategy.

## Introduction

Alzheimer’s disease (AD) is the most common cause of dementia and a progressive neurodegenerative disorder characterized by cognitive decline. Its major pathological features include extracellular β-amyloid (Aβ) plaques, intracellular neurofibrillary tangles composed of pathological tau, gliosis, synaptic loss, and progressive neuronal degeneration(Serrano-Pozo et al., 2011; DeTure and Dickson, 2019).

Synaptic dysfunction and loss occur early and correlate strongly with cognitive decline, making preservation of synaptic integrity an important therapeutic goal. Astrocytes play essential roles in maintaining synaptic function and regulating Aβ metabolism, and their dysfunction may contribute to synaptic degeneration and amyloid pathology in AD. Brain-derived neurotrophic factor (BDNF) signaling through the full-length TrkB receptor (TrkB.FL) promotes neuronal survival, synapse formation, and synaptic plasticity(Huang and Reichardt, 2001; Chao, 2003; Lu et al., 2013; Park and Poo, 2013; Wang et al., 2024). However, BDNF-TrkB signaling is impaired in AD, potentially contributing to synaptic dysfunction, neuronal degeneration, and disease progression(Phillips et al., 1991; Nagahara and Tuszynski, 2011; Lu et al., 2013; Paulo-Ramos et al., 2026). Thus, enhancing endogenous BDNF-TrkB signaling represents a compelling disease-modifying strategy to preserve synapses and neurons in AD.

The mouse *Ntrk2* gene generates three TrkB isoforms through alternative splicing: the full-length receptor (TrkB.FL) and two truncated receptors, TrkB.T1 and TrkB.T2 (collectively referred to as TrkB.T)(Klein et al., 1990; Middlemas et al., 1991; Baxter et al., 1997). TrkB.FL contains an extracellular ligand-binding domain, a single transmembrane domain, and an intracellular tyrosine kinase domain required for canonical BDNF signaling through the PI3K-AKT, MAPK, and PLCγ1 pathways(Berkemeier et al., 1991; Klein et al., 1991; Soppet et al., 1991; Squinto et al., 1991; Klein et al., 1992). In contrast, TrkB.T1 and TrkB.T2 share the same extracellular and transmembrane domains but possess distinct short intracellular tails that lack the kinase domain(Klein et al., 1990; Middlemas et al., 1991). TrkB.T1 accounts for nearly all TrkB.T receptors in the brain (Middlemas et al., 1991; Dorsey et al., 2006) and is expressed predominantly by astrocytes, whereas TrkB.FL is expressed primarily by neurons(Beck et al., 1993; Rose et al., 2003b; Holt et al., 2019; Niu et al., 2024). Accumulating evidence indicates that TrkB.T1 acts as an endogenous negative regulator of BDNF signaling by sequestering BDNF and forming signaling-deficient heterodimers with TrkB.FL(Biffo et al., 1995; Eide et al., 1996; Alderson et al., 2000). TrkB.T1 also regulates astrocyte morphology and function independently of TrkB.FL(Rose et al., 2003b; Fenner, 2012; Holt et al., 2019; Tessarollo and Yanpallewar, 2022). Notably, TrkB.T1 expression is elevated in the brains of patients with AD, raising the possibility that increased TrkB.T1 expression compromises endogenous neuroprotective BDNF-TrkB.FL signaling(Connor et al., 1996; Ferrer et al., 1999; Tessarollo and Yanpallewar, 2022). These observations suggest that reducing TrkB.T expression could enhance endogenous BDNF-TrkB.FL signaling and thereby protect synapses and neurons in AD.

In this study, we investigated whether reducing astrocytic TrkB.T expression ameliorates AD pathology and synaptic loss in 5XFAD mice. We selectively deleted the TrkB-encoding *Ntrk2* gene in astrocytes at 5 weeks of age, resulting in a marked reduction in TrkB.T expression without affecting TrkB.FL levels. Remarkably, astrocytic TrkB.T ablation substantially reduced Aβ plaque burden and prevented synaptic synaptic loss in 5XFAD mice. These findings identify astrocytic TrkB.T as an important regulator of amyloid pathology and synaptic integrity and suggest that reducing TrkB.T expression may represent a promising therapeutic strategy for AD.

## Materials and Methods

### Animals

Mouse strains, Aldh1l1-CreERT2 (IMSR_JAX:031008) (Srinivasan et al., 2016)and 5xFAD (MMRRC_034848-JAX)(Oakley et al., 2006), were obtained from The Jackson Laboratory, and the *Ntrk2^lox/lox^* mouse strain was described previously(Baydyuk et al., 2011). All three mouse strains had been backcrossed onto the C57BL/6J genetic background for at least five generations. We first generated Aldh1l1-CreERT2;*Ntrk2^lox/ox^* and 5xFAD;*Ntrk2^lox/ox^* mice. These mice were then intercrossed, and their offspring were administered tamoxifen (150 mg/kg, i.p.) once daily for eight consecutive days beginning at 5 weeks of age to induce Cre-mediated recombination. Mice were group-housed (4-5 per cage) under standard laboratory conditions with ad libitum access of food and water. Both male and female mice were included in the experiments. All animal procedures were conducted in accordance with institutional guidelines and regulations and were approved by the Institutional Animal Care and Use Committee at the UF Scripps Institute (protocol 16-003-04).

### Immunohistochemistry

All animals were anesthetized with overdose of avertin and perfused transcardially with phosphate buffered saline (PBS) and 4% paraformaldehyde. Brain samples were harvested and postfixed for 24 h in 4% paraformaldehyde, followed by incubation in 30% sucrose solutions until brains sank to the bottom. Brains were sliced into 40-μm sagittal sections a sliding microtome (Leica SM2000R).

For 3,3′-diaminobenzidine (DAB) immunostaining, sections were washed with TBS and quenched with 10% methanol and 3% H_2_O_2_ in TBS for 18 min. After washing with TBS, the sections were blocked in TBS containing 0.4% Triton X-100, 1% glycine, 1% BSA, and 10% normal horse serum for 1 h. Primary antibodies were diluted in blocking buffer and incubated with section at 4°C overnight. Sections were incubated with biotinylated secondary goat anti-rabbit (Vector, BA-1000, 1:200) or goat anti-mouse (Vector, BP-9200-50, 1:200) antibodies for 1 h at room temperature. The VECTASTAIN Elite ABC Kit (PK-6100) were used for ABC reactions following the manufactory’s protocols. After wash with TBS and 100 mM Tris-HCl (pH7.5), sections were developed with 0.05% DAB, 0.003% H_2_O_2_ in 100 mM Tris-HCl (pH7.5) for 3-5 min. After transferring onto Superfrost plus slides, sections were dehydrated by sequential incubation in 70% ethanol (5 min), 95% ethanol (5 min), 100% ethanol (twice, 5 min each), and xylene (twice, 10 min each). DPX Mountant and coverslip were applied.

For fluorescent immunohistochemistry, sections were washed in PBS and then blocked in PBS with 10% normal horse serum and 0.3% Triton X-100 for 1 h. Sections were then incubated at 4°C overnight with primary antibodies diluted in PBS containing 10% normal horse serum and 0.3% Trion X-100. Secondary antibodies were diluted in the same diluent as primary antibodies and incubated at room temperature for 1 h. Sections were incubated with DAPI (2.5 μg/ml) in PBS for 5 min. Sections were mounted onto Superfrost plus slides, and Fluoro Gel mounting medium (Electron microscope science, 17985-10) was applied.

The following primary antibodies were used for the immunohistochemistry: chicken anti-GFAP (1:500; encore), guinea pig anti-Iba1 (1:2000), mouse anti-amyloid β (1:500; Biolegend), guinea pig anti-bassoon (1:1000; Synaptic Systems #141 004), mouse anti-PSD-95 (1:100; Synaptic Systems # 124 011). Fluorescent images were captured using a Nikon C2+ confocal microscope or a Leica TCS SP8 microscope.

### Density and size of astrocytes

Every 6^th^ sagittal brain sections were used for quantification of astrocytic density and size. DAB-stained sections were examined under a Nikon Eclipse E800 microscope. Stereo Investigator software (MBF Bioscience) was used to quantify the density and size of GFAP-immunoreactive astrocytes in different brain regions.

### Western blotting

Prefrontal cortical tissues were dissected on ice using a coronal Brain Matrix (Roboz, SA-2175). Dissected brain tissues were lysed on ice for 30 min in lysis buffer containing 1mM EDTA, 50mM Tris-HCl, 150mM NaCl, 1% NP-40, 0.5% sodium deoxycholate, 0.1% sodium dodecyl sulfate, and freshly added protease inhibitors (Roche Complete Protease Mini, #4693159001). Lysates were centrifuged at 15,000Å∼g for 30 min at 4 °C, and supernatants were saved. Protein samples were run on SDS-PAGE gels and transferred to PVDF membrane. Membrane was blocked with Odyssey Blocking Buffer (Thermo Fisher Scientific). The following antibodies were used: goat anti-TrkB (1:1000; R&D systems), Mouse anti-α tubulin (1:5000; sigma Aldrich). Appropriate IRDye infrared secondary antibodies (LI-COR Biosciences) were used at a dilution of 1:10,000. Odyssey Infrared Imaging System (Image Studio Lite Ver 4.0, LI-COR Biosciences) was used to detect and quantify signals of target proteins.

#### Epitope-preserving magnified analysis of the proteome (eMAP)

The eMAP protocol was adapted and optimized from Park et al (Joha Park and Qiangge Zhang, 2021). Briefly, 12-month-old mice were deeply anesthetized with an overdose of avertin and transcardially perfused with 1x PBS, followed by 4% paraformaldehyde (PFA). Brains were dissected and post-fixed in 4% PFA for 48 h at 4⁰C. Samples were washed in PBS containing 0.02% sodium azide (PBSN) for 24 h at 4⁰C, followed by additional 24 h at room temperature. Brains were trimmed to 3-mm blocks and incubated in hydrogel monomer solution for 72 h at 4⁰C, protected from light. For hydrogel embedding, samples were placed individually in cassettes containing eMAP solution (30% acrylamide, 0.1% bisacrylamide, 10% sodium acrylate, and 0.03% VA-044 initiator in 1x PBS). Polymerization was performed in a 37⁰C water bath under a continuous flow of nitrogen gas at 10 psi for 3-4 h. Following polymerization, samples were incubated overnight in PBSN at 37⁰C. Embedded tissue blocks were sectioned at a thickness of 100 µm using a vibratome (Leica VT1200) with the following settings: frequency, 60 Hz; amplitude, 1.2 mm; and feed rate, 0.8 mm/s.

Sections were cleared overnight at 37⁰C with shaking in clearing solution containing 6% sodium dodecyl sulfate, 0.1 M phosphate buffer, 50 mM sodium sulfite, and 0.2% sodium azide (pH7.4) to remove lipids. Sections were then denatured in the clearing solution at 95⁰C for 10 min and washed in 1x PBS containing 0.1% Triton X-100 and 0.02% sodium azide before immunostaining.

### Statistical Analysis

Statistical analyses were performed using GraphPad Prism software. Statistical significance was defined as p < 0.05. The statistical tests, sample sizes, and significance thresholds are specified in the corresponding figure legends.

## Results

### Astrocyte-specific ablation of TrkB.T expression in 5xFAD mice

5xFAD mice co-express amyloid precursor protein (APP) and presenilin 1 (PS1) carrying five familial AD (FAD) mutations under the control of the *Thy1* promoter(Oakley et al., 2006). These mice develop robust Aβ plaques, microgliosis, and astrogliosis by 2 months of age and exhibit synaptic loss by 9 months of age(Oakley et al., 2006; de Pins et al., 2019; Forner et al., 2021). To investigate whether reducing astrocytic TrkB.T expression ameliorates AD pathology, we induced astrocyte-specific *Ntrk2* deletion in 5xFAD mice beginning at 5 weeks of age, before the onset of overt pathology.

Because TrkB.T is predominantly expressed in astrocytes, which express relatively little TrkB.FL(Beck et al., 1993; Rose et al., 2003b; Holt et al., 2019; Niu et al., 2024), we reasoned that astrocytic-specific *Ntrk2* deletion would markedly reduce TrkB.T levels while preserving TrkB.FL expression. To test this hypothesis, we crossed astrocyte-specific, tamoxifen-inducible *Aldh1l1-CreERT2* mice with *Ntrk2^lox/lox^*mice to generate *Aldh1l1-CreERT2*;*Ntrk2^lox/lox^* mice, which were subsequently crossed with 5xFAD;*Ntrk2^lox/lox^* mice. All offspring were administered tamoxifen daily for eight consecutive days beginning at 5 weeks of age to induce Cre-mediated recombination. Four experimental genotypes were examined: *Ntrk2^lox/lox^* (control), 5xFAD;*Ntrk2^lox/lox^* (5xFAD), *Aldh1l1-CreERT2*;*Ntrk2^lox/lox^* (astrocyte-specific *Ntrk2* conditional knockout; aNtrk2-cKO), and 5xFAD;*Aldh1l1-CreERT2*;*Ntrk2^lox/lox^*(5xFAD;aNtrk2-cKO).

To determine the effects of astrocytic *Ntrk2* deletion on TrkB expression, we performed immunoblot analysis of cortical lysates from the four genotypes (Fig. 1A). TrkB.T levels were reduced by ≥80% in both aNtrk2-ckO and 5xFAD;aNtrk2-cKO mice compared with control and 5xFAD mice (Fig. 1B). In contrast, TrkB.FL levels were comparable among the four genotypes (Fig. 1C). These results demonstrate that astrocyte-specific *Ntrk2* deletion markedly reduces TrkB.T expression in the brain while preserving TrkB.FL levels, establishing a genetic model for investigating the role of astrocytic TrkB.T in AD pathology.

**Figure 1.**
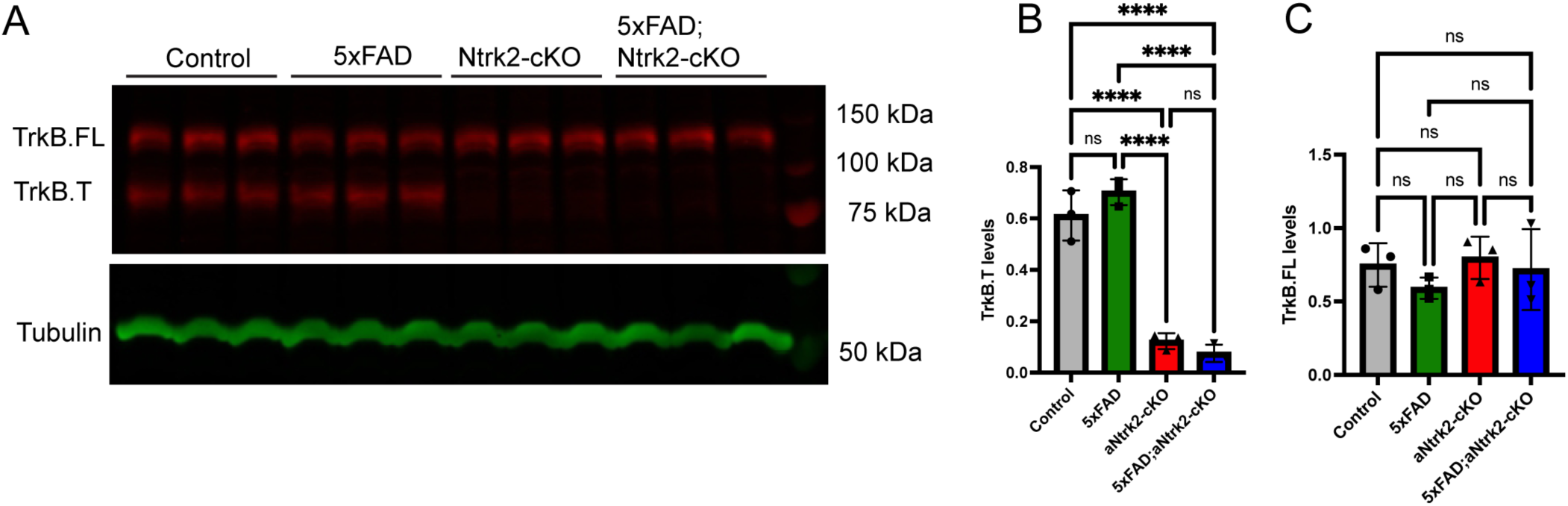
Astrocytic Ntrk2 deletion reduces cortical TrkB.T levels without affecting TrkB.FL expression. (**A**) Immunoblots of TrkB.FL (upper band) and TrkB.T (lower band) in cortical lysates from 1-year-old control, 5xFAD, aNtrk2-cKO, and 5xFAD;aNtrk2-cKO mice. *^a^*-tubulin was used as a loading control. (**B**, **C**) Quantification of TrkB.T (B) and TrkB.FL (C) levels normalized to *^a^*-tubulin. Data are presented as mean ± SEM (n = 3 mice per group). Statistical significance was determined by one-way ANOVA followed by Tukey’s multiple comparisons test: ns, not significant; ****p<0.0001.

### Astrocytic TrkB.T ablation does not alter gliosis in 5xFAD mice

Neuroinflammation is a hallmark of AD brain and is characterized by the accumulation of reactive astrocytes and activated microglia surrounding amyloid plaques(Akiyama et al., 2000). TrkB.T has been shown to mediate intracellular Ca^2+^ transients and regulate cytoskeletal dynamics through interactions with RhoGDI, an inhibitor of Rho family GTPases(Rose et al., 2003a; Ohira et al., 2005; Ohira et al., 2006). Thus, astrocytic TrkB.T ablation could alter astrocyte reactivity in response to Aβ deposition and potentially influence microglial activation. To investigate this possibility, we performed immunohistochemistry for glial fibrillary acidic protein (GFAP) and ionized calcium binding adaptor molecule 1 (Iba1) to assess astrogliosis and microgliosis, respectively, using brain tissue from mice at 6 months of age. Reactive astrocytes typically exhibit increased GFAP expression and cellular hypertrophy(Escartin et al., 2021), whereas microgliosis is associated with increased Iba1 immunoreactivity, changes in microglial morphology, and, in some contexts, increased microglial density(Kettenmann et al., 2011).

Consistent with previous findings (Oakley et al., 2006), GFAP immunohistochemistry revealed increased numbers of GFAP-positive astrocytes and pronounced astrocytic hypertrophy in the somatosensory cortex (SSC; Fig. 2A) and hippocampal CA1 region (Fig. 2F) of 5xFAD mice compared with control mice. To quantify these changes, we used Stereo Investigator and Neurolucida software to assess the density of GFAP-positive astrocytes, the number and length of their processes, and soma size in both regions.

**Figure 2.**
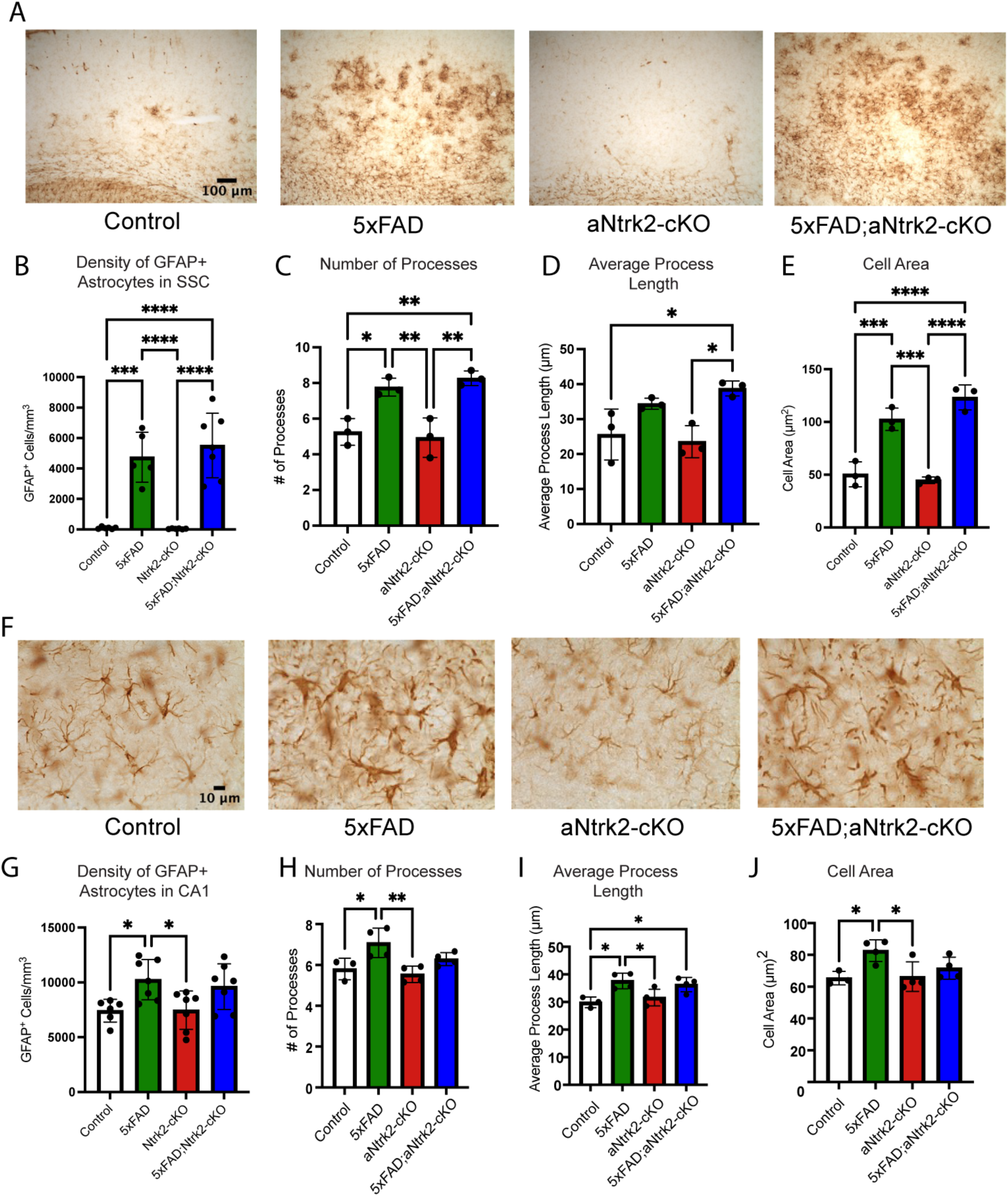
Astrocytic TrkB.T ablation does not alter astrocyte density or morphology in 5xFAD mice. (**A**) Representative images of GFAP immunohistochemistry in the SSC of 6-month-old control, 5xFAD, aNtrk2-cKO, and 5xFAD;aNtrk2-cKO mice. (**B**–**E**) Quantification of GFAP-positive astrocyte density and morphological parameters in the SSC. (**F**) Representative images of GFAP immunohistochemistry in the hippocampal CA1 region of 6-month-old mice. (**G**–**J**) Quantification of GFAP-positive astrocyte density and morphologic parameters in CA1. GFAP immunohistochemistry was performed on every sixth section. Astrocyte density was quantified in four sections per mouse, and 15 astrocytes per mouse were analyzed for morphological measurements (n = 3-7 mice per group). Statistical significance was determined by one-way ANOVA followed by Tukey’s multiple comparisons test: *p<0.05, **p<0.01, ***p<0.001, and ****p<0.0001.

Quantitative analysis revealed a higher density of GFAP-positive astrocytes in 5xFAD mice than in control mice (Fig. 2B, G), particularly in the SSC, where basal GFAP expression is low (Fig. 2A). Astrocytes in 5xFAD mice also exhibited marked hypertrophy, characterized by increased process number (Fig. 2C, H), process length (Fig. 2D, I), and soma size (Fig. 2E, J). However, none of these parameters differed significantly between control and aNtrk2-cKO mice or between 5xFAD and 5xFAD;aNtrk2-cKO mice. These findings indicate that astrocytic TrkB.T ablation does not alter basal astrocyte morphology or GFAP-defined reactive astrogliosis in 5xFAD mice.

Activated microglia typically exhibit enlarged somata, fewer and thicker processes, and increased Iba1 immunoreactivity, resulting in a greater area occupied by Iba1 staining. Consistent with microglial activation, immunofluorescence analysis revealed that the area occupied by Iba1 immunoreactivity was markedly increased in the hippocampal CA1 region and SSC of 5xFAD mice compared with control mice (Fig. 3). However, astrocytic TrkB.T ablation did not significantly affect this increase in either region (Fig. 3). Together with the GFAP immunohistochemistry results (Fig. 2), these findings indicate that astrocytic TrkB.T ablation does not alter overt astrogliosis or microgliosis in 5xFAD mice, as assessed by GFAP and Iba1 immunoreactivity, respectively.

**Figure 3.**
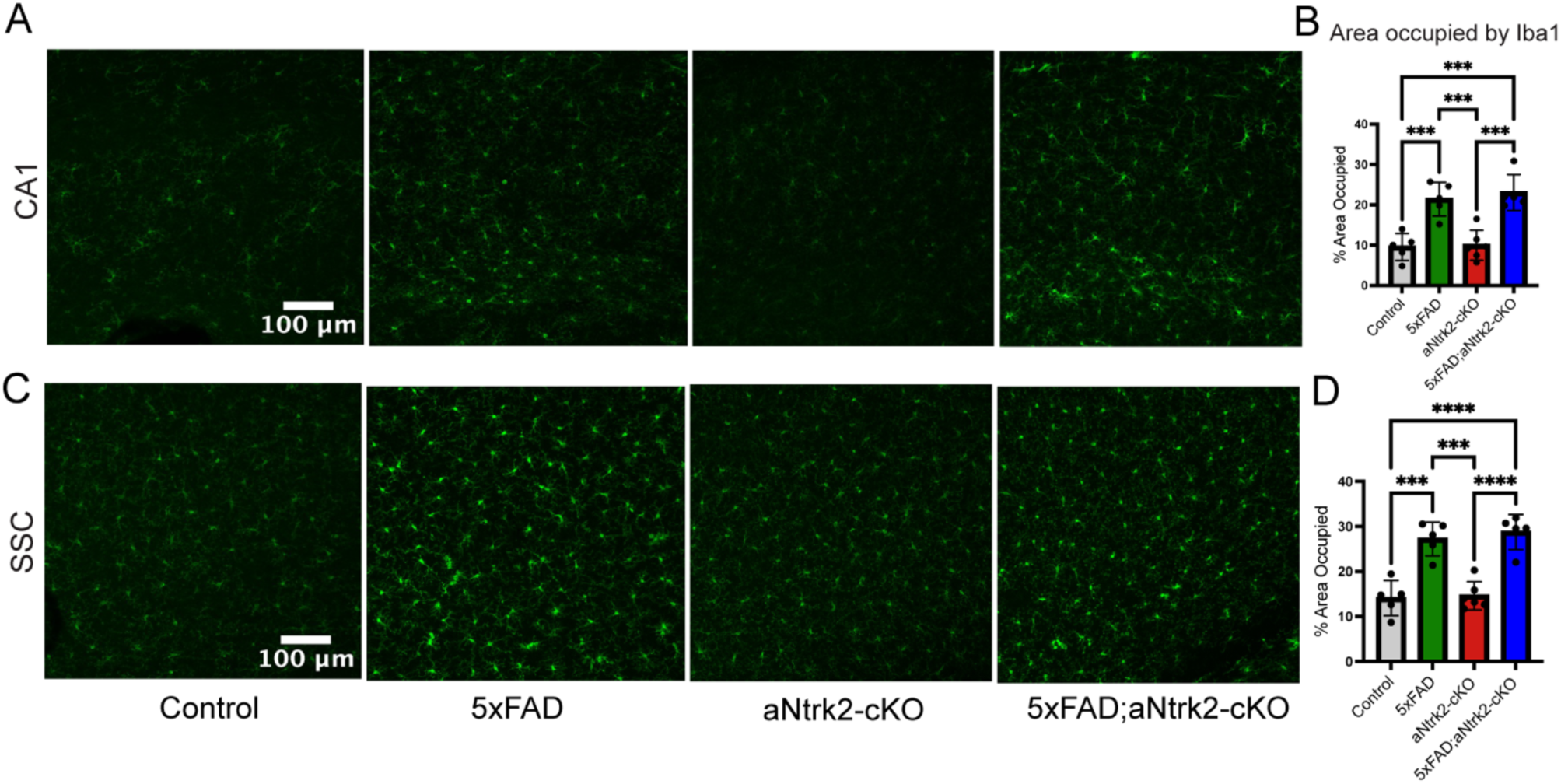
Astrocytic TrkB.T ablation does not alter microgliosis in 5xFAD mice. (**A**, **C**) Representative images of Iba1 immunofluorescence in the hippocampal CA1 region (A) and SSC (C) of 6-month-old control, 5xFAD, aNtrk2-cKO, and 5xFAD;aNtrk2-cKO mice. (**B**, **D**) Quantification of the area occupied by Iba1 immunoreactivity in the CA1 (B) and SSC (D). Iba1 immunohistochemistry was performed on every sixth section, and the area occupied by Iba1 immunoreactivity was quantified in four sections per mouse (n = 5-6 mice per group). Statistical significance was determined by one-way ANOVA followed by Tukey’s multiple comparisons test: ***p<0.001 and****p<0.0001.

### Astrocytic TrkB.T ablation markedly reduces Aβ plaque burden

Astrocytic TrkB.T ablation could influence Aβ accumulation by altering APP processing through enhanced BDNF-TrkB.FL signaling and/or promoting Aβ clearance through changes in astrocyte function. To determine whether astrocytic TrkB.T ablation affects amyloid pathology, we performed Aβ immunohistochemistry on sagittal brain sections from 6-month-old control, 5xFAD, aNtrk2-cKO, and 5xFAD;aNtrk2-cKO mice. As expected, numerous Aβ plaques were detected in the cortex and hippocampus of 5xFAD mice but were absent in control and aNtrk2-cKO mice (Fig. 4A). Remarkably, astrocytic *Ntrk2* deletion markedly reduced Aβ plaque accumulation in 5xFAD mice (Fig. 4A).

**Figure 4.**
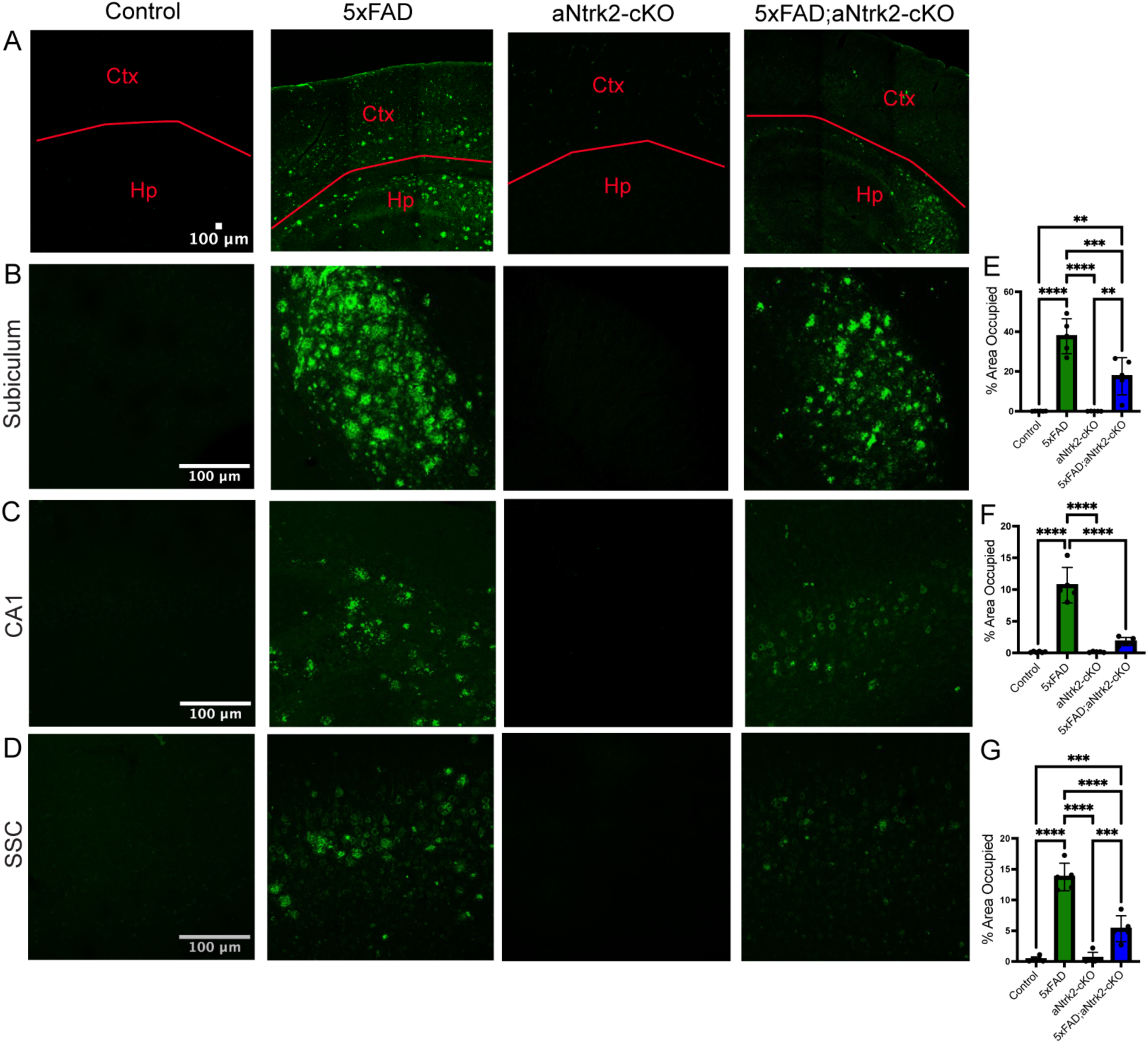
Astrocytic TrkB.T ablation reduces Aβ plaque burden in 5xFAD mice. (**A**) Representative images of Aβ immunofluorescence in the cortex (Ctx) and hippocampus (Hp) of control, 5xFAD, aNtrk2-cKO, and 5xFAD;aNtrk2-cKO mice at 6 months of age. (**B**–**D**) Representative images of Aβ immunofluorescence in the subiculum (B), CA1 (C), and SSC (D) of the four genotypes. (**E**–**G**) Quantification of the area occupied by Aβ immunoreactivity in the subiculum (E), CA1 (F), and SSC (G). Aβ immunohistochemistry was performed on every sixth section, and the area occupied by Aβ immunoreactivity was quantified in four sections per mouse (n = 5-6 mice per group). Statistical significance was determined by one-way ANOVA followed by Tukey’s multiple comparisons test: **p<0.01, ***p<0.001, and ****p<0.0001.

To quantify this effect, we measure the percentage of tissue area occupied by Aβ immunoreactivity in the subiculum, hippocampal CA1 region, and SSC. Consistent with previous reports(Oakley et al., 2006), Aβ deposition was particularly prominent in the subiculum of 5xFAD mice (Fig. 4B-D). Astrocytic TrkB.T ablation reduced Aβ plaque burden by 53% in the subiculum (Fig. 4B, E), 83% in CA1 (Fig. 4C, F), and 61% in the SSC (Fig. 4D, G). Thus, astrocytic TrkB.T ablation markedly reduces Aβ plaque burden across multiple brain regions in 5xFAD mice, with reductions ranging from 53% to 83%. The magnitude of this reduction is comparable to the substantial decreases in amyloid plaque burden reported following anti-Aβ immunotherapy in preclinical AD models(Da Mesquita et al., 2021). These findings identify astrocytic TrkB.T as an important regulator of Aβ accumulation and suggest that its reduction may represent a promising strategy for attenuating amyloid pathology in AD.

### Astrocytic TrkB ablation preserves synaptic integrity in 5xFAD mice

To determine whether astrocytic TrkB.T ablation protects against synaptic degeneration in 5xFAD mice, we employed epitope-preserving Magnified Analysis of Proteome (eMAP) (Park et al., 2021)to examine synaptic structures in the brains of 1-year-old control, 5xFAD, aNtrk2-cKO, and 5xFAD;aNtrk2-cKO mice. eMAP enables high-resolution imaging of synaptic structures using conventional confocal microscopy and facilitates their three-dimensional (3D) reconstruction. We performed immunofluorescence staining for Bassoon and PSD-95 to label presynaptic active zones and postsynaptic densities (PSDs), respectively. Confocal z-stacks were acquired from the SSC, and Imaris software was used to reconstruct and quantify Bassoon- and PSD-95-positive puncta in 3D (Fig. 5A).

**Figure 5.**
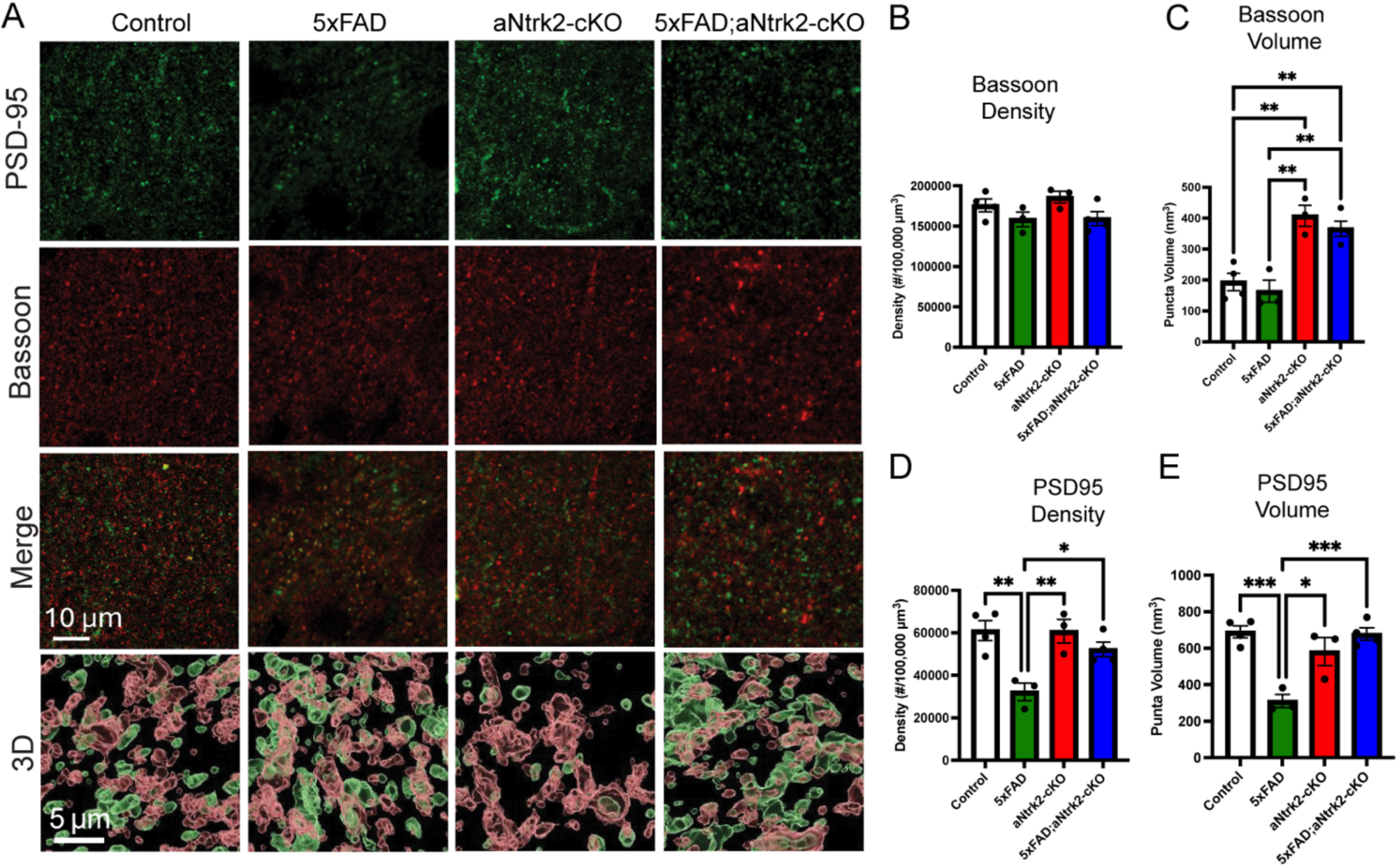
Astrocytic TrkB.T ablation preserves postsynaptic structures in 5xFAD mice. (**A**) Representative single-plane confocal images of PSD-95 and 3D reconstructions of Bassoon and PSD-95 immunofluorescence in the SSC of 1-year-old control, 5xFAD, aNtrk2-cKO, and 5xFAD;aNtrk2-cKO mice. (**B**, **C**) Quantification of the density (B) and volume (C) of Bassoon-positive puncta. (**D**, **E**) Quantification of the density (D) and volume (E) of PSD-95-positive puncta. Four confocal z-stacks per mouse were used for 3D reconstruction and quantification of Bassoon- and PSD-95-positive puncta (n = 3-4 mice per group). Statistical significance was determined by one-way ANOVA followed by Tukey’s multiple comparisons test: *p<0.05, **p<0.01, and ***p<0.001.

Quantitative analysis revealed that astrocytic TrkB.T ablation markedly increased the size of Bassoon-positive puncta without affecting their density (Fig. 5B, C). This finding suggests that reducing astrocytic TrkB.T expression promotes enlargement of presynaptic active zones, potentially through enhanced BDNF-TrkB.FL signaling. Importantly, astrocytic TrkB.T ablation prevented the reductions in both the size and density of PSD-95-positive puncta observed in 5xFAD mice (Fig. 5D, E), indicating that TrkB.T ablation protects against AD-associated loss of postsynaptic structures. Together, these findings demonstrate that astrocytic TrkB.T ablation preserves synaptic structural integrity in 5xFAD mice and suggest that it may also enhance presynaptic function by promoting active zone enlargement.

## Discussion

This study demonstrates that selective *Ntrk2* deletion in astrocytes markedly reduces the expression of TrkB.T, which negatively regulates BDNF-TrkB.FL signaling, while preserving TrkB.FL expression. This selective reduction could enhance endogenous neuroprotective BDNF-TrkB.FL signaling. Consistent with the established role of BDNF-TrkB.FL signaling in synaptic maintenance and plasticity (Lu et al., 2013; Park and Poo, 2013), we found that astrocytic TrkB.T ablation prevented reductions in the density and size of postsynaptic densities in 5xFAD mice while increasing the size of presynaptic active zones. Remarkably, astrocytic TrkB.T ablation also substantially reduced Aβ plaque burden across multiple brain regions. Together, these findings suggest that reducing astrocytic TrkB.T expression could provide multiple therapeutic benefits by enhancing endogenous neuroprotective signaling, attenuating amyloid pathology, and preserving synaptic integrity, potentially improving synaptic function. Thus, targeting astrocytic TrkB.T represents a promising therapeutic strategy for AD that could simultaneously address amyloid pathology and synaptic degeneration.

Enhancing BDNF–TrkB signaling has long been considered a promising therapeutic strategy for AD. BDNF gene delivery to the entorhinal cortex was shown to reverse synaptic loss and improve cognitive performance in an APP-transgenic mouse model without altering amyloid plaque burden (Nagahara et al., 2009). Furthermore, treatment with a TrkB agonistic antibody preserved synaptic structures and ameliorated cognitive deficits in APP/PS1 mice (Wang et al., 2020). Several studies have also reported that treatment with putative small-molecule TrkB agonists, including 7,8-dihydroxyflavone and its derivatives, improves cognitive performance in 5xFAD mice (Devi and Ohno, 2012; Aytan et al., 2018; Chen et al., 2021). However, subsequent studies have questioned whether these compounds directly activate TrkB, complicating the interpretation of their therapeutic mechanisms (Boltaev et al., 2017; Pankiewicz et al., 2021). In contrast to approaches relying on exogenous BDNF or receptor agonists, our study identifies an alternative strategy to potentially enhance endogenous BDNF-TrkB.FL signaling by reducing the expression of its inhibitory truncated receptor. Remarkably, astrocytic TrkB.T ablation not only prevented AD-associated reductions in the density and size of postsynaptic densities but also increased the size of presynaptic active zone, revealing an additional and potentially important synaptic benefit. Although the functional consequences of active zone enlargement remain to be determined electrophysiologically, this unexpected finding raises the possibility that astrocytic TrkB.T ablation could not only preserve synaptic integrity but also enhance neurotransmitter release at surviving synapses. Thus, reducing astrocytic TrkB.T expression may offer a therapeutic strategy that combines synaptic protection with potential enhancement of synaptic function, ultimately helping to ameliorate cognitive deficits in AD.

One striking finding of this study is the marked reduction in Aβ plaque burden across multiple brain regions of 5xFAD;aNtrk2-cKO mice compared with 5xFAD mice. Treatment with putative small-molecule TrkB agonist 7,8-dihydroxyflavone and its derivatives has been reported to reduce Aβ plaque burden in AD mouse models (Devi and Ohno, 2012; Aytan et al., 2018; Chen et al., 2021). However, enhancing BDNF-TrkB signaling does not invariably reduce amyloid pathology. BDNF gene delivery and long-term administration of a specific, high-affinity TrkB agonistic antibody improved cognitive performance without affecting Aβ plaque burden in AD mouse models (Nagahara et al., 2009; Wang et al., 2020). Conversely, heterozygous deletion of *Ntrk2* in 5xFAD mice exacerbated memory deficits without altering Aβ plaque burden. These observations suggest that the profound reduction in Aβ plaque burden following astrocytic TrkB.T ablation may involve mechanisms beyond neuronal BDNF-TrkB.FL signaling. Moreover, given the uncertainty surrounding the direct activation of TrkB by 7,8-dihydroxyflavone and its derivatives (Boltaev et al., 2017; Pankiewicz et al., 2021), their reported effects on amyloid pathology may involve TrkB-independent mechanisms. One possibility is that astrocytic TrkB.T regulates Aβ accumulation through its effects on astrocyte function. Under pathological conditions, TrkB.T signaling in astrocytes may impair Aβ clearance, either directly by affecting astrocytic uptake and degradation of Aβ or indirectly by modulating microglial clearance mechanisms. Alternatively, astrocytic TrkB ablation may reduce neuronal Aβ production by altering APP processing. Determining whether astrocytic TrkB.T ablation reduces Aβ production, enhances its clearance, or both will be an important direction for future studies and may reveal previously unrecognized mechanisms by which astrocytes regulate amyloid pathology in AD.

## Acknowledgements

This work was supported by the grants from the National Institutes of Health to BX (R01 DK134650) and HVM (F31 NS143305). YC is the Len and Norma Klorfine graduate fellow at the Herbert Wertheim UF Scripps Institute for Biomedical Innovation & Technology.

